# Fine Endmesolithic Fish Caviar Meal Discovered by Proteomics of Foodcrusts

**DOI:** 10.1101/332882

**Authors:** Anna Shevchenko, Andrea Schuhmann, Günter Wetzel

## Abstract

The role of aquatic resources in ancient economies and paleodiet is important for understanding the evolution of prehistorical societies. However, conventional archaeometric approaches lack organismal specificity, are affected by abundant environmental contaminants and do not elucidate food processing recipes. We performed proteomics analysis of charred organic deposits adhered on early ceramics from Mesolithic-Neolithic inland site Friesack 4 (Brandenburg, Germany). Proteomics of foodcrust from a vessel attributed to Endmesolithic pottery identified fine carp roe meal and revealed details of a prehistorical culinary recipe. We propose that Endmesolithic occupants of Friesack at the end of the 5^th^ millennium BC utilized fish as a food reserve and adopted delayed-return subsistence strategy. These data contribute to better understanding of the dietary context of Neolithic transition in European inland.

## Introduction

The major inducements underlying developments in prehistorical societies were access and use of natural resources, which also played a key role in Neolithic transition across Europe. The role of aquatic resources during this critical phase in European prehistory remains in the focus of longstanding debates particularly regarding inland regions [1-6].

Inland prehistorical localities were often situated close to rivers or lakeshores. Although it is conceivable that fish and water plants were a part of the diet of their inhabitants their general subsistence strategies did not necessarily rely on freshwater resources [7]. Our understanding of the role of aquatic resources in economy of prehistorical aceramic communities relies on assemblage of recovered artefacts: zooarchaeological materials (shellfish, bones, scales), catching or processing tools and related art objects - zoomorphic figurines, paintings, adornments. Additionally, analysis of DNA recovered from ancient fishbones was used for identification of fish species [8]. Systematic carbon and nitrogen isotopic analysis of associated human remains (bones, teeth or hairs) shed a light on dietary importance of aquatic products in the population [9, 10]. However, the related archaeological artefacts are absent at many sites. Fish bones and scales are not well preserved in acidic soils and are often underrepresented in archaeofaunal assemblages if no environmental sampling or sieving was applied. Isotopic composition of human remains is influenced by many factors including physiology of the individual [11,12]. Apart of it, analysis of stable isotopes is significantly limited in the regions where diverse dietary resources are available and did not applicable for estimation of dietary contribution of fresh water products at inland sites [7].

Neolithic transition in European inland was accompanied by appearance and spread of early ceramics which serialization is important for chronological reconstruction. Charred food remains (or foodcrusts) often found on these early pottery is a valuable molecular evidence, particularly for evaluation of the dietary role of aquatic resources in prehistorical communities. Isotopic composition and pattern of small organic molecules in ceramic foodcrusts - lipids, fatty acids, sterols and a few known molecular biomarkers, - enable distinguishing terrestrial, marine and fresh water materials (reviewed in [13,14]). However, they do not provide species-specific information and reveal very little details about cooking recipe. Also employed analytical procedures are sensitive to contemporary contaminates.

Analysis of proteins in archaeological organic residues is advantageous for the interpretation of their complex composition (reviewed in [15]) and are essential also for ceramic foodcrusts. Ancient proteins carry aging-specific chemical modification which helps distinguishing them from modern proteneous contaminants [16,17]. Protein sequences provide direct information about the species of origin [18-20]. Furthermore, their biological properties (enzymatic activity or organismal localization) could elucidate the type of applied processing technologies [21,22]. Thus, identification of protein composition in foodcrusts might also assist reconstruction of cooking recipe of prehistorical foods.

In this work we present proteomics analysis of foodcrusts from ceramics recovered at the inland Mesolithic-Neolithic site Friesack 4 (Brandenburg, Germany). The site is known for its complex stratigraphy and excellent preservation of numerous predominantly Mesolithic artefacts in the over 2m deep cultural occupation layer. This unique collection also comprises early ceramics including Friesack-Boberger group [23,24] which represents the most southern example of Mesolithic pottery in the Northern European lowland. A small fraction of the pottery fragments from Friesack preserves visible foodcrusts. Their protein compositions shed light on prehistorical cooking practice, paleodiet and help understanding the process of transformation from hunter-gatherer groups to sedentary society in European inland.

## Material and Methods

### Geographical location and history of exploitation of Friesack 4

Friesack 4 is a Mesolithic-Neolithic bog site near a small town Friesack (52° 43′ 59″ N, 12° 34′ 59″ E) in Havelland administrative district (Brandenburg, Germany) ca 70km west-northwest from Berlin (Fig 1A). It was discovered by amateur archaeologist Max Schneider at the beginning of the 20^th^ century when amelioration and draining bogs along Rhin River revealed first artefacts [25], and later explored by several expeditions leaded by H. Reinerth, S. Wenzel and particularly B. Gramsch (Fig 1A-D).

Friesack 4 is situated in the Warsaw-Berlin ice margin valley on the shore of the Rhin River. Between Preboreal and Atlantic the landscape around the site presented afforested lowland along the water boundary interjected by ground moraines. According to archaeobotanical, palynological and ichthyoarchaeological analyses, the region was first grown over with mixed pine and birch forest which was later replaced by deciduous trees, and was inhabited by broad variety of wild animals [26-29]. From as early as the end of the 10^th^ millennium BC groups of Mesolithic hunter-gatherers frequented at the Friesack site which was located at the shore of a lake [26]. Depositional events at the site suggested that till the end of Mesolithic in the middle of the 6^th^ Millennium BC it was visited 50-60 times [30]. Elevation of water level in terminal Boreal and Atlantic caused partial inundation and formation of massive fens around Friesack 4 and also probably resulted in a short occupational pause at the site [31]. Despite significantly changed landscape the territory remained attractive for hunting and fishing in terminal Mesolithic and was later occasionally visited by Neolithic groups settled in the Friesack vicinity [23]. The site was utilized exclusively as a seasonal hunting station occupied between early spring and late fall [32] rather than a year-around settlement. Trace amount of pollen detected at Friesack 4 [26] suggested that the place was not used as a cropland. Most likely Neolithic settlers cultivate the land south or west from Friesack 4.

**Fig 1.**
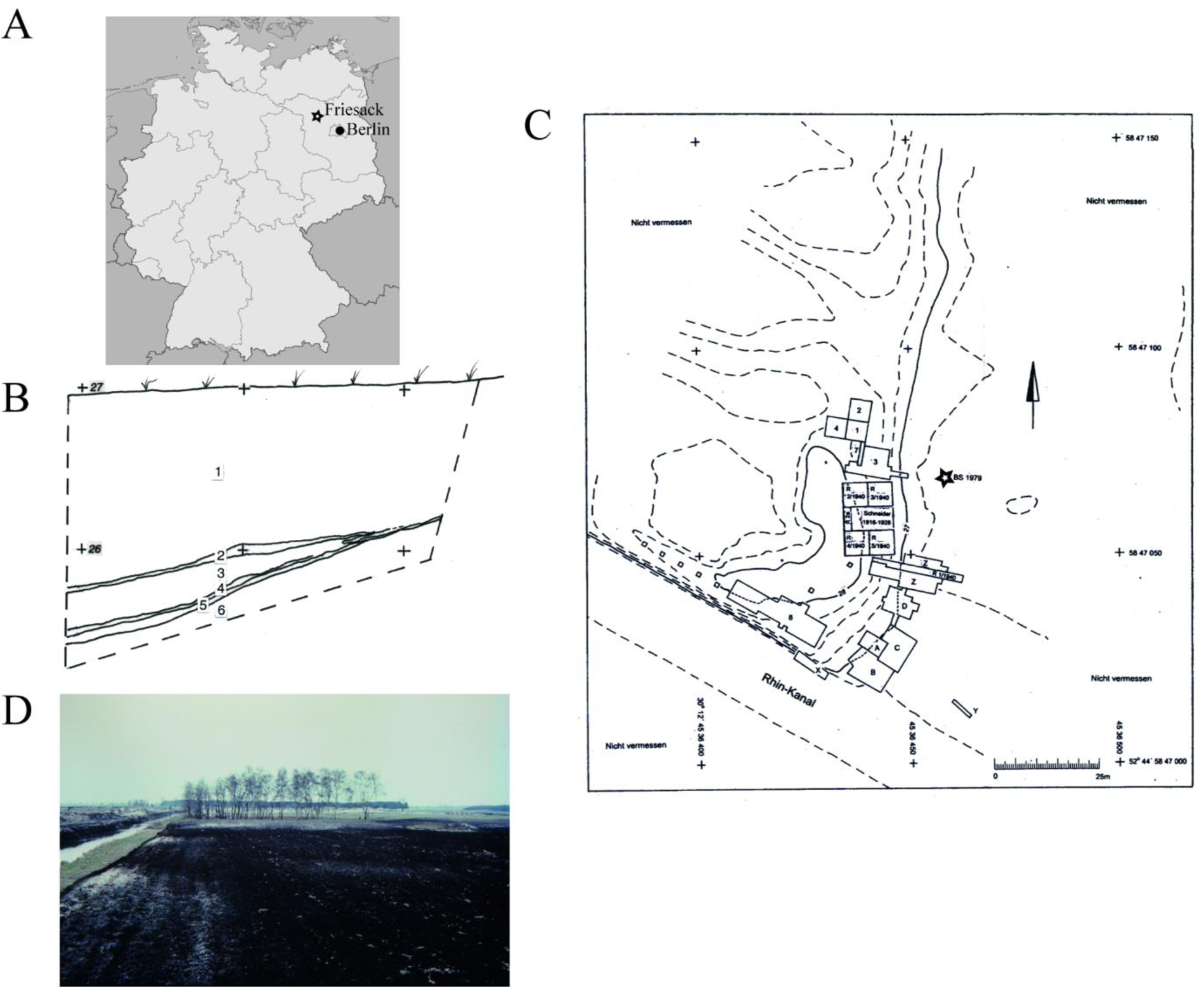
Archaeological site Friesack 4. **A:** Location in Havelland administrative district, Brandenburg, Germany. **B:** Layout of archaeological excavations undertaken at the Friesack 4 site by M. Schneider (1916–1928), H. Reinerth (R 1-5/1940), B. Gramsch (1977–1989; 1998; 1-4, 7, A-D, X-Z) and S. Wenzel (2000–2001; 8). The position of the test trench 1979 is indicated by asterisk (graphic rendering the original map from Bodo Hildebrand (†) and Dieter Becker). **C.** South profile of the test trench 1979: 1 - peat; 2, 4 – yellowish grey light sand; 3, 5 – organic silt; 6 – Pleistocene sand (graphic rendering the original drawing from B. Gramsch). D. A view to the South profile of the Friesack 4 site after the upper layers of peat and sand were ploughed; with the permission of B. Gramsch.

### Friesack 4 collection of archaeological artefacts

Friesack 4 collection of artefacts comprises more than 150000 objects, the vast majority of which are Mesolithic. Apart of numerous lithics, it also includes objects made from organic materials: bone, wood, bast, pitch, antler - which have been well preserved in the natural environment of peat bog interspaced with sandy layers [27]. The collection encompasses nearly all groups of artefacts which were manufactured by Mesolithic occupants: from arrowheads and bast nets to decorated carapace of freshwater turtle and birch bark tar chewing gum – and shows substantial similarity with Maglemose culture [33,34]. Ceramic fragments represent a very small fraction of the collection. No prehistorical burials were found in the closed vicinity apart of remains of two individuals briefly mentioned in the excavation reports from 40^th^, and a few single fragments of reported at the site [27,35].

Whereas the Mesolithic history of Friesack 4 has been intensively studied (Gramsch, Kloss, Teichert, Görsdorf, Benecke, Jahns, Schmölcke and others) little is known about later phases of exploitation of the site. Only in a few recent years, its less abundant Neolithic artefacts including unique samples of early ceramics awakened interest of archaeologists [23,24,32].

### Archaeological samples

**#3258** (Inv. No 1977:7/3258 T): a group of 12 small to middle-size shards of the same fragment of a coarse undecorated pottery. The group was recovered from the lowest occupational layer (#4) of the south profile of a test trench made in the refuse zone northeast of the settlement by B.Gramsch in 1979 (Landesmuseum für Ur- und Frühgeschichte Potsdam, Germany) (Fig 1B,C). The foodcrusts were adhered at inner surface of basal and rim shards. Individual shards were glued together shortly before sampling organic deposit. Radiocarbon dating, typology and reconstruction of the vessel profile are presented in this work.

**#3251.1** (Inv. No 1977:7/3251.1): a body shard of round-bodied vessel decorated with three drop-shaped imprints. It was recovered closely to the position of the #3258 from the layer #5 of the same trench in 1979 (Fig 1B). Charred organic crust on the internal surface was radiocarbon dated 3946–3776 calBC (ARR 15048). The fragment was attributed to Early Neolith, Brzesc Kujawski group (?) ([23] Abb.6.6; [24] Abb.6.2).

**#3157.1** (Inv. No 1977:7/3157.1): a body fragment of massive S-shaped vessel with all-over regular fingernail impressions. It was recovered from unseparated Neolithic occupation layer underlying humus, in area 3 quadrat A15 in 1984 (Fig 1C). Foodcrust adhered on inner surface was radiocarbon dated 4336-4241 calBC (ARR 15046). The shard was attributed to the Friesack-Boberger group ([23] Abb. 6.7; [24] Abb.6.1).

**#3078 (**Inv. No 1977:7/3078): a rim shard of a funnelbeaker decorated with three irregularly placed angular impressions. The shard was recovered from humus mixed cultural layer of quadrat D5 in 1983 (Fig 1C). Charred foodcrust on the internal surface was radiocarbon dated 3628-3376 calBC (ARR 15044) ([23] Abb.6.1; [24] Abb.7.3).

The shards were gently cleaned with a brush to remove loose soil contaminants, wrapped in paper and stored in carton boxes at the Brandenburger Office for Cultural Heritage Preservation (Brandenburgisches Landesamt für Denkmalpflege und Archäologisches Landesmuseum, Wünsdorf, Germany).

### Modern reference samples

Fresh carp roe was purchased at the fish farm Heinz Mueck Fischhandel GmbH, Dresden; fish muscle tissue (*S. salar from aquatic culture, Norway*) obtained by local vendor in Dresden. For preparation of fish broth, a portion of ca 125g of fish muscle tissue was cooked for 30min in ca 300ml of salty water on low hit in an open pot. A sample of in-house manufactured pottery glue was provided for analysis by the workshop of the Brandenburger Archaeological Museum in Wünsdorf, Germany.

### Radiocarbon dating of #3258

^14^C dating was carried out at the Poznań Radiocarbon Laboratory, Poznań, Poland (Poz-80609) leaded by Prof. T. Goslar. The data were analysed with OxCal software v.4.2.4 [36] and calibrated using intCal13 atmospheric curve [37].

### Proteomics analysis

20-35mg of encrusted organic deposits from internal surface of #3251.1, #3157.1 and #3078 shards and from the base of #3258 were scratched off using cleaned scalpel and spatula. Each foodcrust was independently sampled twice except for #3251.1 (due to lack of available material). The material was first crashed with a disposable pestle into fine powder in an Eppendorf tube. Then proteins were extracted with 65 mM Tris HCl buffer (pH 6.8) containing 2% sodium dodecylsulfate (SDS) and loaded together with the slurry on a pre-cast 1 mm 12% polyacrylamide gel (BioRad Laboratories, Munich, Germany). After electrophoretic separation the gel was visualized by Coomassie staining, sample gel lane cut into 10 slices each of which was then process individually. Each gel slab was cut into 1×1mm cubes, washed twice with 50% acetonitrile in 50mM ammonium bicarbonate and dehydrated with neat acetonitrile. Then the gel pieces were rehydrated in 10mM ammonium bicarbonate containing 10% of acetonitrile and 12ng/µl trypsin (modified, sequencing grade, Promega, Mannheim) and their protein content overnight *in-gel* digested at 37°C. The resulting peptide mixtures were extracted twice with exchange of 5% formic acid and 100% acetonitrile, the extracts pulled together and dried down. Then the peptides were re-suspended in 20µl of 5% formic acid and 5µl were taken for LC-MS/MS analysis.

Proteins from modern reference samples and a portion (20mg) of in-house made adhesive for pottery reconstruction were extracted, processed and analysed by proteomics on the same way. The list of protein identified in ten blank LC-MS/MS runs (*in-gel* digests of void gel slabs) was used to sort out typical protein contaminants introduced by sample preparation and handling.

Peptides recovered from each gel slab were separately analysed by LC-MS/MS on an Ultimate3000 nano-UPLC system interfaced on-line to a LTQ Orbitrap Velos hybrid tandem mass spectrometer (Thermo Fisher Scientific, Bremen, Germany) as described in. The UPLC system was equipped with Acclam PepMap^tm^ 100 75 µm x 2cm trapping column and 75µm x 15cm separating column packed with 3 µm diameter C18 particles (Thermo Fischer Scientific, Bremen). Peptides recovered from each gel slab were separated using 180min linear gradient (solvent A – 0.1% formic acid in water, solvent B – 0.1% formic acid in neat acetonitrile). Velos instrument was operating in CID mode where 1 MS scan (R=60000) was followed by MSMS acquired from top 20 most abundant precursor ions. Normalized collision energy was set on 35 and dynamic exclusion time on 25 s. The lock-mass function was set to recalibrate MS1 scans using the background ion (Si(CH_3_)_2_O)6 at m/z 445.1206. The spectra were converted into mgf format (Mascot Generic Format) and combined into a single file. Database search was performed by Mascot software v.2.2.04 (Matrix Sciences Ltd, London, UK) against all species or *Actinopterygii* (for fish reference samples) protein sequences in NCBI protein database (April 2015, 64057457 entries) under the following settings: mass accuracy 5ppm and 0.5Da for precursor and fragment ion respectively; enzyme specificity – trypsin; maximal number of allowed miscleavages – two; variable modifications – methionine and proline oxidized; N-terminal protein acetylation; cysteine propionamide; asparagine and glutamine deamidated [16,17]. Protein hits were accepted if matched by minimum two unique peptides satisfying the following stringent criteria: peptide sequences were longer than seven amino acid residues, peptide ion scores exceeded the MASCOT homology threshold and was above the value of 30. The result of the database search for fish reference samples was evaluated by Scaffold software (Proteome Software, Portland, v.4.7.5) using 90% peptide and protein probability thresholds and minimal of g two matched peptides. Calculated False Discovery Rates (decoy FDR) for peptides and proteins were below 0.1%. Chromatographic peaks of selected peptides were integrated using Xcalibur software (Thermo Fischer Scientific) and confirmed by the corresponding fragmentation spectra. Deamidation rate of asparagine and glutamine residues for individual peptides was calculated as ratio of integrals of XIC of their modified and non-modified form.

Where specified, sequence similarity search for peptides identified by proteomics was performed by MS BLAST program (http://genetics.bwh.harvard.edu/msblast/index.html) [38] against NRDB95 database (v.2016_06); protein BLAST was performed on https://www.ncbi.nlm.nih.gov/ against NCBI protein database.

### Scanning Electron Microscopy (SEM)

Uncoated dry samples were fixed on a metal holder and imaged by SEM on a Magellan 400 FEG-SEM instrument (FEI, Eindhoven, The Netherlands) in immersion mode with accelerating voltages of 3 and 5 kV, beam current 100 pA under variable magnification.

## Results

### Morphological analysis, age and typology of the vessel #1977:7/3258 T

First, it was necessarily to define the age and typology of the selected pottery fragments. Attribution and ^14^C radiocarbon analysis of ceramics #3251.1, #3157.1 and #3078 were reported previously [23,24] whereas the pottery #3258 was not analysed. #3258 comprised 12 middle-size to small potshards originating from same vessel which was broken during excavation. The fragments included well preserved lip and basal shards and allowed reliable reconstruction of the pottery profile.

**Fig 2.**
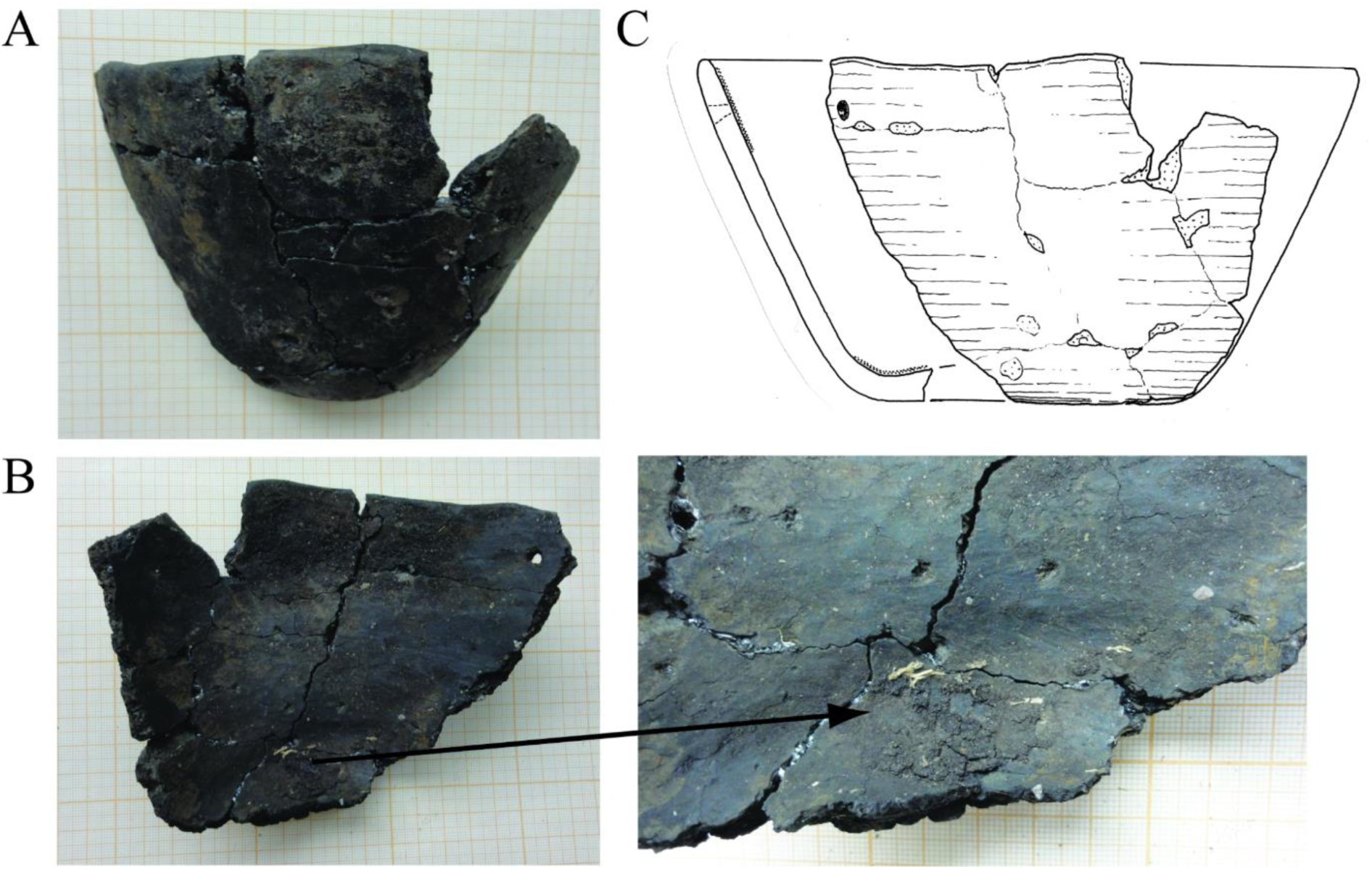
Fragment of endmesolithic pottery #1977:7/3258 T from Friesack 4. **A:** side view; **B:** inner view, secondary conical puncture is seen at the upper right corner. Panel on the right shows basal fragment of the dish with foodcrust. **C:** Graphical reconstruction of the profile of the bowl #1977:7/3258 T.

The 10.5cm high vessel with 20cm round opening had straight rim and rounded lip, conically ascended wall and rounded flat base with 9.0cm diameter (Fig 2A,B). The pottery was made of medium coarse fired clay tempered with grit and had well smoothed, unglazed, dark brown surface. The body and basal fragment of the vessel are remarkably massive – 0.7cm at the top and 0.6 at the bottom with 1cm-thick base getting slightly stronger toward the middle. One of the rim shards had secondary conical puncture with 0.5cm outer and 0.3cm inner diameter located ca 1.5cm under the edge of the lip. This presumably mending hole was drilled from outside inwardly after pottery burning (Fig 2C). Small amount of dry organic material was adhered as a narrow strip on the inner surface along the rim and as a few millimetre thick dark brown deposit with cracked texture on the base inside of the vessel #3258.

^14^C radiocarbon analysis of the crust from the rim of the #3258 resulted in a broad range between 4300 and 4000 calBC (Supplementary Fig S1). Considering that freshwater reservoir effect might also significantly alter radiocarbon dating [13] and that its offset is not established for the site of excavation, to aid the result we looked at other artefacts recovered in the close proximity to #3258. Two potshards - #3256.1 attributed to the Endmesolithic Friesack-Boberger group [23] (Supplementary Fig S2) and #3251.1 assigned to Early Neolith – were found in layers N2 and N5 of the same trench respectively (Fig 1C). These age estimates concurred with ^14^C dating of #3258 and positioned the sample in the time span between Endmesolithic and Early Neolith. The combination of unusually massive walls, rounded base and rim of the undecorated coarse conical vessel #3258 did not occur in Neolithic pottery in the Brandenburg area. Pottery with flat base and conical walls appeared in the region in Rössener culture (site Dyrotz 37 [24,39]) which is relatively isochronal with Friesack 4 finding, though rounded base, rounded rim and the thickness of walls are not typical for Rössener ceramics. Considering radiocarbon dating, pottery specific characteristics and additionally the fact that the potshards #3258 were found in the lowest occupational layer of the trench, we attributed #3258 to the Endmesolithic pottery of the recently described Friesack-Boberger group [23,24]. Friesack-Boberger group has certain similarity with Ertebølle ceramics which was replaced by Funnelbeaker pottery at around 4000BC [40] – the time point which also coincides with the #3258 dating.

### Protein composition of foodcrusts from Friesack 4 pottery

Proteomics identified from 70 to 300 proteins of different organismal origin in foodcrusts from Friesack 4 pottery (Supplementary dataset S1): human, microbial, plant, insect, terrestrial animals, and fish. To facilitate the interpretation of the result, proteins considered as a common background were removed from the list of hits and remaining proteins were grouped accordingly to their organismal origin (Table 1, Supplementary dataset S1). Next, we assessed deamidation rates of asparagine und glutamine residues in matched peptides. Non-enzymatic deamidation of these amino acids is aging-related process which could be used for distinguishing ancient from modern proteins [16,17,19,41]. While in the fresh samples observed deamidation rate of these two residues is <1%, the conversion could be almost completed in proteins over 3800 years old [16].

**Table 1.** Overview of Friesack 4 foodcrusts.

| N | Sample name | Archaeological inventory N | <sup>14</sup>C dating |  | Ceramics typology | Proteomics analysis result (number of identified proteins)<sup>1</sup> |  |  |
| --- | --- | --- | --- | --- | --- | --- | --- | --- |
|  |  |  | BP | calBC |  | Common protein background<sup>2,3</sup> | Contemporary<sup>3</sup> environmental contaminants | Ancient proteins<sup>3</sup> |
| 1 | #3258 | 1977:7/3258 T | 5350±30 | 4300-4050 | Friesack-Boberger | Human proteins (32); Hen egg ovalbumin; Bovine milk proteins (1) | Bacteria: *Streptomyces* spp (177), other (7) Fungi<sup>3</sup> (6) | Fish vitellogenins and parvalbumins |
| 2 | #3251.1 | 1977:7/3251.1 | 5042±25 | 3946-3776 | Brzesc Kujawski (?) [23,24] | Human proteins and multispecies hits (90); Bovine milk proteins (4) | Bacteria: *Streptomyces* spp (169); other (27) Fungi: *Fusarium oxysporum* (11); other (12) Plants: *Cucurbita maxima*, phloem (2); *Betula pendula*, pollen (1) | Collagen, *Suidae* (2) |
| 3 | #3157.1 | 1977:7/3157.1 | 5419±27 | 4336-4241 | Friesack-Boberger [23,24] | Human proteins (34); Bovine milk proteins (2) | Bacteria: *Streptomyces* spp (8); other (2); Fungi (13) |  |
| 4 | #3078 | 1977:7/3078 | 4705±22 | 3628-3376 | Funnelbeaker [24] | Human proteins (42); *Picea sitchensis* (2); Bovine milk proteins (2); Bovine collagen (2) | Insect proteins (5) Plants: *Triticeae* spp, seed (5); Fam. *Lamiaceae*; green (1) |  |

<sup>1</sup> – full list of identified proteins is shown in the Supplementary Table S1.
<sup>2</sup> – protein background includes human skin-associated, hair and saliva proteins observed in multiple independent GeLC analyses (data not shown) , laboratory standards and proteins detected in pottery glue sample (Supplementary Table S1).
<sup>3</sup> - the assignment is based on deamidation rate of glutamine and asparagine residues in detected peptides. Proteins with residual deamidation rate below 1%, as also observed in model protein samples, were considered contemporary.

### Protein background in Friesack 4 samples

Common protein background comprised human skin, saliva and hair proteins introduced during sample handling [16,21,22], conserved multispecies matching proteins which organismal origin cannot be determined, e.g. histones or actin, and laboratory standards (hen egg ovalbumin) [42]. Because the fragments of the vessel #3258 T were glued together before sampling the foodcrust, we also analysed a sample of the home made adhesive and included identified proteins in the background list (Supplementary dataset S1). Interestingly, minute amount of silk protein sericin (gi 112984400, *B.mori*) which is widely used as moisturizing agent in modern skin and hair cosmetics (up to 20% w/w in some cream formulations) [43] was identified among other contaminants.

Bovine milk proteins were detected in all Friesack foodcrusts and also in the sample of pottery adhesive (Table 1). The peptide YLGYLEQLLR from bovine alpha-S1-casein being present in each of samples. Glutamine residue (Q) of this peptide in 3500-years old dairy samples was completely deamidated [16,22]. However, this and other glutamine and asparagine residues in milk proteins from Friesack foodcrusts remained unmodified and therefore these proteins were considered as modern contaminants.

Microbial proteins: the bulk of remaining proteins (except for crust #3078) has microbial origin (Table 1, Supplementary dataset S1) and includes species typical for the natural microorganismal community of a peat bog. The majority of microbial proteins matched to *Streptomyces* spp.- the bacteria which is common to humid soils and might be involved in lysis of cellulose [44,45]. Other detected microbial proteins belong to bacteria inhabiting meromictic freshwater lakes as well as fungal proteins from plant-associated genera *Fusarium* and from plant pathogens occurring in microbiome of peat bog [46]. Though, no *Aspergillus* spp which are a common abundant compound of destructive microflora on archaeological artefacts [47], were detected. Asparagine and glutamine residues in microbial proteins of Friesack samples were not deamidated.

Plant proteins were detected in crusts #3078 and #3251. They originated from seeds of a plant belonging to *Triticeae* family, green of a plant from *Lamiaceae* family, phloem of squash *Cucurbita maxima* and birch pollen (Table 1, Supplementary dataset S1). Although they could be associated with foods, particularly seeds from *Triticeae* plant, none of asparagine and glutamine residues in matching peptides were deamidated. Several insect muscle proteins matching various species were identified in crust #3078. They were also not deamidated and presumably belong to environmental contaminants.

### Modern and ancient collagens in Friesack 4 foodcrusts

Two core collagens - alpha1(I) and alpha2(I) - matching altogether 25 peptides were identified in foodcrusts #3078 and #3251.1 and, in trace amounts, also in the control sample (pottery adhesive) (Tables 1, 2). Whereas the majority of collagen-matched peptides was identical to multiple animal collagens, a few of them in #3078 and control sample were unique for protein entries from *Bovidae* (acc no P02453 and P02465) and in crust #3251.1 - from pig (acc no A0A1S7J210 and A0A1S7J1Y9). Notably, asparagine and glutamine residues in these eight collagen peptides were partially deamidated only in crust sample 3251.1. Composition, species specificity and deamidation rate of detected peptides (Table 2) suggested that foodcrusts #3078 and #3251.1 comprised modern *Bovidae* collagen as a contaminant. In addition, the foodcrust #3251.1 also includes minute amount of pig collagen, which most likely of an ancient origin.

**Table 2.**
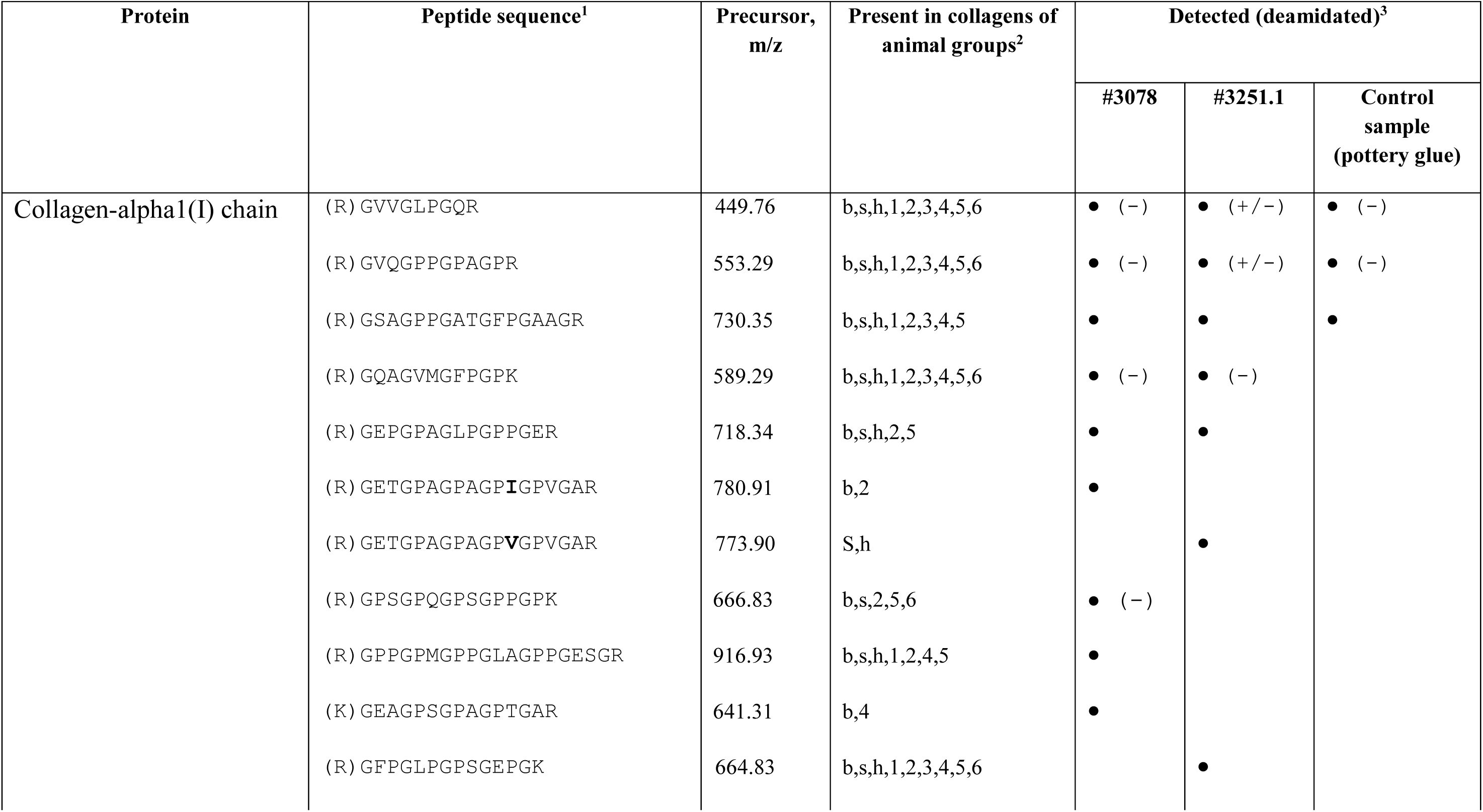
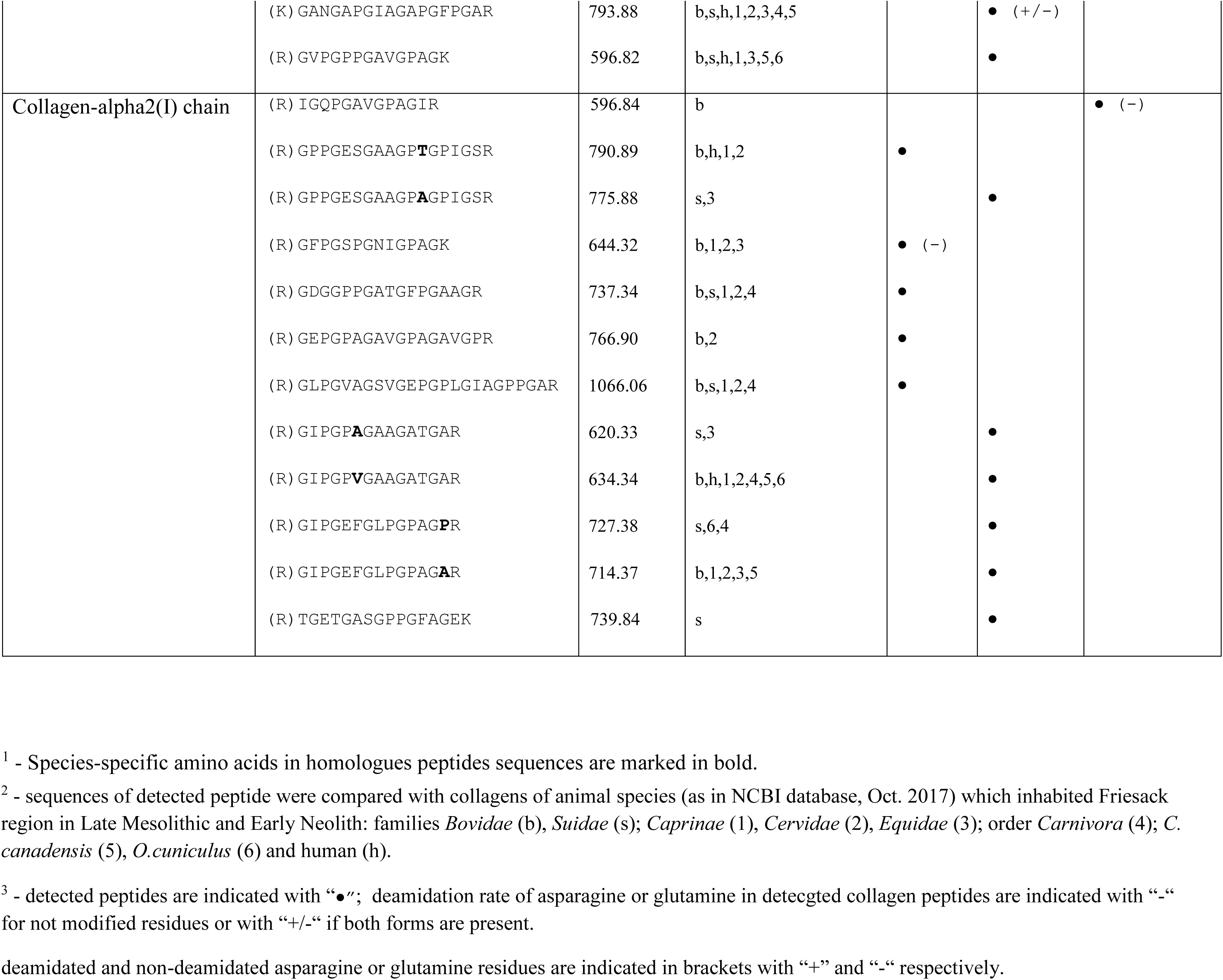
List of peptides matched to collagens in foodcrusts #3078 and #3251.1.

### Ancient fish proteins in the foodcrust #3258, their properties and species attribution

Two fish proteins, vitellogenin and parvalbumin, matched with seven and three peptides, respectively, were only identified in the fooodcrust from the base shard of the #3258 vessel (Tables 1 and 3). Four out of seven vitellogenine peptides were unique for *Cypriniformes* order including common carp *Cyprinus carpio* and the remaining three were present in multiple vitellogenin sequences from various *Actinopterygii* (Table 3). Detected parvalbumin peptides belong to a conserved part of the sequence and are present in many fish species. MS BLAST sequence similarity search matched them all together to a single sequence from common carp (*Cypriniformes*) as the top hit (Table 3, Fig 3). The results suggested that the foodcrust #3258 comprised material from a *Cypriniformes* specie that concurred with the site-specific fish population dominated in Neolith [32].

**Table 3.** List of peptides matched to fish vitellogenin and parvalbumin in foodcrust #3258*.

| N | Sequence of peptides detected in #3258 | Gene Identifier of the matched protein sequence<sup>a</sup> | Organismal order | Deamidation rate of asparagine and glutamine residues (%)<sup>b</sup> |  |
| --- | --- | --- | --- | --- | --- |
|  |  |  |  | #3258 | Fresh carp roe |
|  |  | *Vitellogenin* |  |  |  |
| 1 | VQVDAILALR | 399761434, 4572552, , 15778562, 156713467, 124518427, 556825564, 157419156, 284097898, 332712555, 662036448, 831290949 | *Cypriniformes, Clupeiformes*<br>*Characiformes* | + (100%) | + (<1%) |
| 2 | LELEVQVGPR | 399761434, 4572552, , 15778562, 742165772, 108863148, 556825564, 157419156, 284097898, 332712555 | *Cypriniformes, Esociformes* | + (88%) | + (<1%) |
| 3 | AYLAGAAADVLEIGVR | 399761434, 4572552, 15778562, 124518427, 15778562, 157419156, 284097898, 662036448 | *Cypriniformes* | + | + |
| 4 | AEAGVLGEFPAAR | 399761434, 4572552, 15778562, 156713467, 108863148, 556825564, 284097898, 63100501, 662036448 | *Cypriniformes* | + | + |
| 5 | EIV<u>ML</u>GYGXXXXK<sup>c</sup> | 742165772, 1024933123, 662036450, 4572552, 157419156, 399761434, 124518427, 160333705, 1025281290 | *Cypriniformes, Esociformes* | + | EVV<u>ML</u>GYGSMIAR<sup>c</sup> |
| 6 | GI<u>IN</u>LLQL<u>N</u>VK | 15778562, 742165772, 5725516, 499021581, 554825686, 86143840, 156530049 | *Cypriniformes, Esociformes, Cichliformes* | + (89%) | GILN<u>IL</u>QL<u>N</u>LK<sup>c</sup> (<3%) |
| 7 | LPIIVTTYAK | 156713467, 691424740, 1020581238 | *Cypriniformes* | + | LPITVTTYAK<sup>c</sup> |
| *Parvalbumin* |  |  |  |  |  |
| 1 | AADSF<u>N</u>YK | 966641035d | *Various Actinopterygii (including Cypriniformes)* | + (85%) | + |
| 2 | SGFIEEDELK | 966641035d | *Various Actinopterygii (including Cypriniformes)* | + | + |
| 3 | LFL<u>Q</u>NFSAGAR | 966641035d | *Various Actinopterygii (including Cypriniformes)* | + (100%) | + (not deamidated) |
| 4 | LFL<u>Q</u>NFSGSAR | 966641035d | *Various Actinopterygii (including Cypriniformes)* | + (100%) | n/a |

\* - data are combined from two analyses. The table includes only peptides satisfying the criteria described in Materials and Methods.
Underlined amino acids were detected modified
<sup>a</sup> - sequences of these proteins comprise peptide which are identical to those detected in #3258 or its homologues resulted from single nucleotide substitution (as for example valine to isoleucine replacement).
<sup>b</sup> - calculated as ratio of integrals of the corresponding peptide chromatographic peaks. “+” – stays for peptides detected in the sample; “n/a.” – no peptide detected.
<sup>c</sup> – matches to several partially overlapping fragmentation spectra acquired from precursors with different mass.
<sup>d</sup> – accession number of the best MS BLAST match

**Fig 3.**
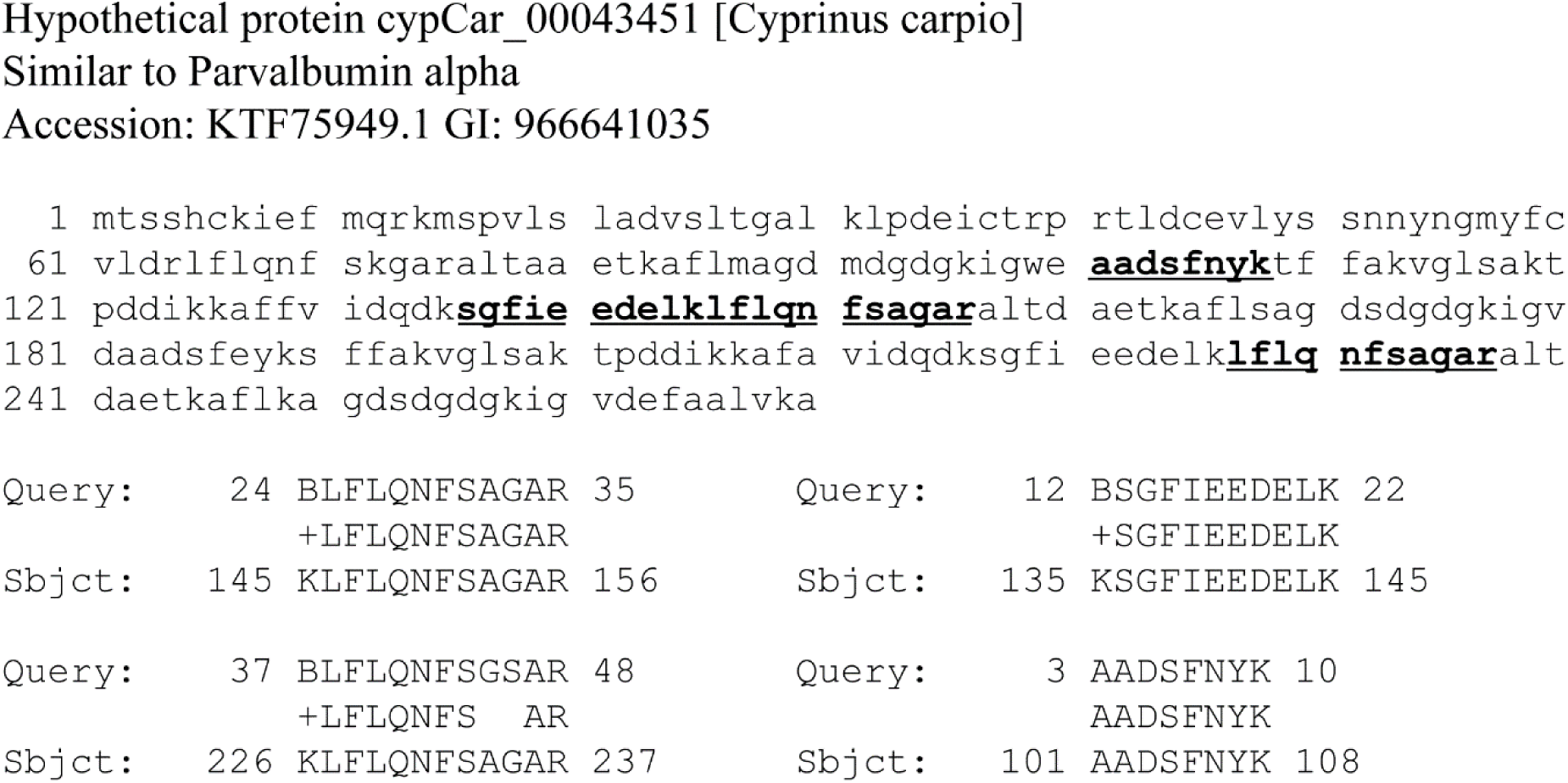
Parvalbumin peptides detected in #3258 basal crust are matched to *C. carpio* sequence by MS BLAST. “Query” includes sequences of peptides identified in the sample; “Sbjct” - sequences of corresponding peptides matched to *C.carpio* protein (acc no KTF75949.1).

Vitellogenin is a major proteins in fish egg yolk [48] thus one of ingredient of the meal in #3258 could be fish caviar. Parvalbumins are small proteins which are abundant in white (non-fatty) muscle of fishes where they presumably promote fibre relaxation [49]. However, no proteins which build the bulk of muscle (myosin, tropomyosin or myoglobin) have been identified in the #3258 residue. To find out whether parvalbumin is detectable also in fish roe we performed proteomic analysis of fresh carp roe sac (unseparated eggs and skin) (Supplementary dataset S1). As anticipated, vitellogenins were the most abundant components of roe matching more than 40% of totally over 65 000 spectra; parvalbumins were present in trace amounts among more than 1800 identified proteins (Supplementary dataset S1). Parvalbumins possess remarkable chemical and thermal stability [50] and we assumed that, together with highly abundant vitellogenin, they are remainders of what once were *Cypriniformes* roe sac.

In contrast to other proteins identified in Friesack samples, all asparagine and glutamine residues in peptides matched to vitellogenin and parvalbumin were almost completely deamidated (Fig 4, Table 3). To prove that deamidation of these specialized proteins does not naturally occur in living fish and is not influenced by thermal treatment, we analysed a sample of fresh carp roe (Table 3, Supplementary dataset S1) and fish muscles proteins after cooking in salty broth (data not shown). In both instances glutamine and asparagine residues were not affected. The deamidation ratio in fish proteins from the crust #3258 was concordant with previously reported for archaeological samples of comparable age [16] and with predicted half-life of unmodified asparagine and glutamine residues [41]. It confirmed the ancient origin of protein material and supported the attribution of #3258 sample.

**Fig 4.**
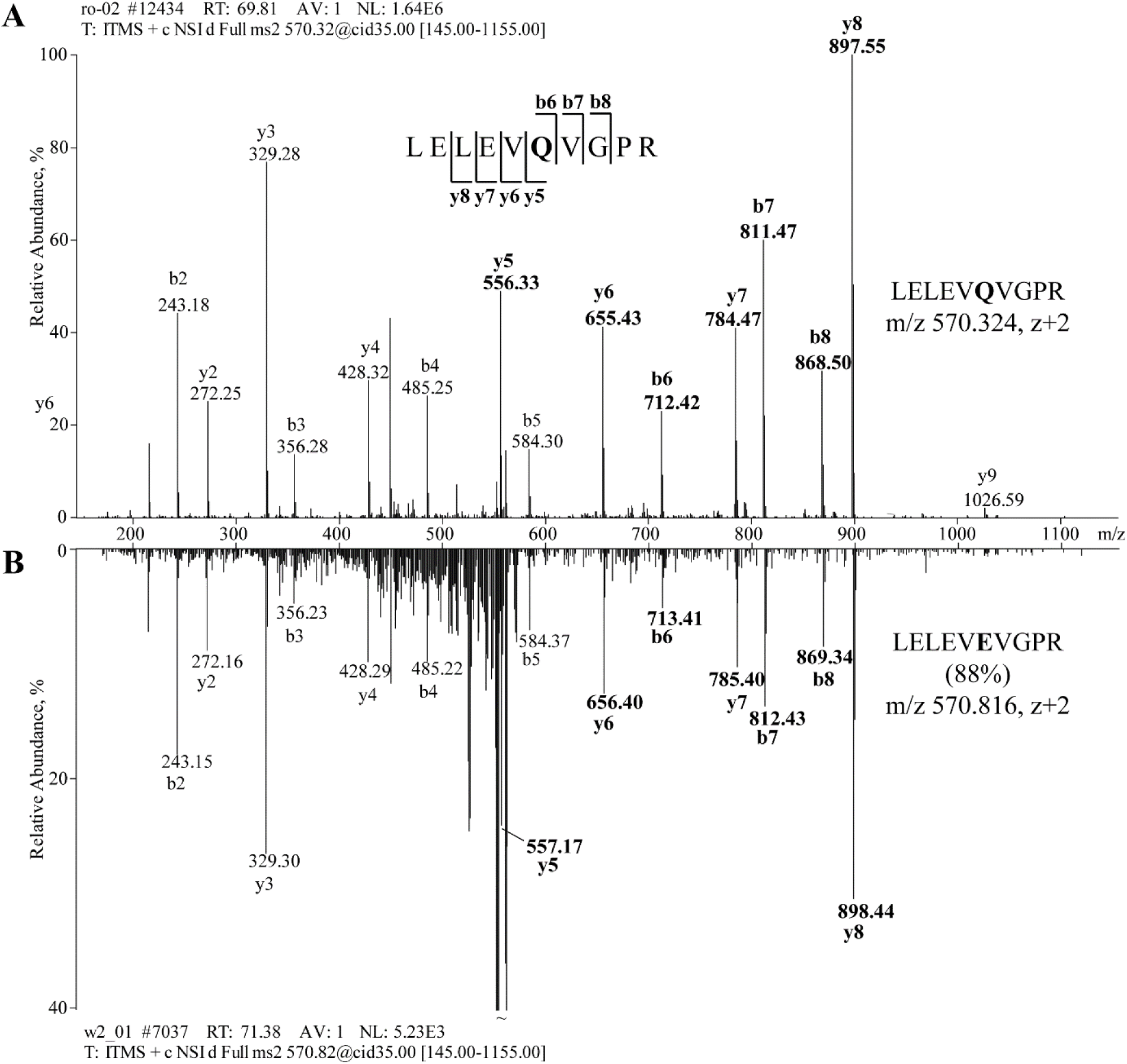
Abundant deamidation of the glutamine residue (Q) in fish vitellogenin from foodcrust #3258. Upper panel: Fragmentation spectrum of vitellogenin peptide LELEV**Q**VGPR (m/z 570.32, z +2) identified in fresh carp roe. In foodcrust #3258, 88% of glutamine residue in this peptide was converted into glutamic acid (E); mirrored image of fragmentation spectrum of peptide LELEV**E**VGPR detected in #3258 is shown at the lower panel. Masses of fragment ions including Q or E at the position six are shown in bold.

### Electron microscopy of foodcrusts from pottery #3258

Electron microscopy of rim and basal foodcrusts from #3258 pottery (Fig 5) did not detect fish microremains although microscopic fish bones and scales are reported in archaeological foodcrusts [51-53]. It corroborates with our assumption that only roe and not the whole fish was cooked in this pottery.

Rim and basal crusts had different texture and density. Rim deposit comprises plant microremains - broken microstructure of xylem, phloem and vascular cuticles of leaves and stems (Fig 5A-E). At some positions, the surface of the rim residue was covered with a dense layer of what might be Streptomyces-like bacterial spores with “fluffy” ornamentation, and ca 1µm salt microcrystals. Two typical conifer bisaccate pollens were incrusted into rim deposit. In contrast, the dark brown basal crust had dense and homogeneous texture with a few light-yellowish grains of sand (Fig 5F). Basal charred food remains were pressed against sand when the broken pottery was thrown away and landed upside down on sandy supralittoral in the refuse zone of the site.

**Fig 5.**
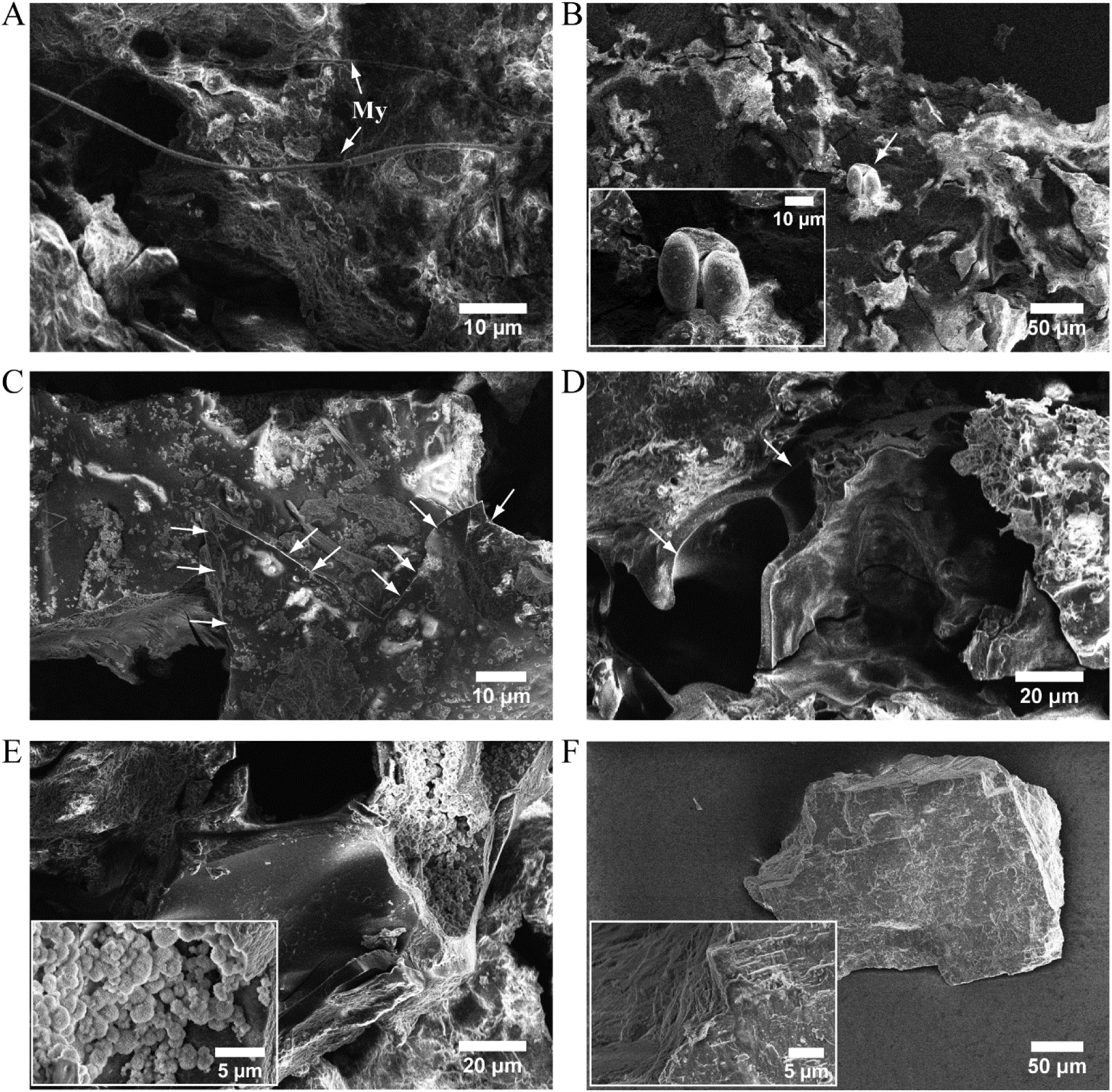
Electron microscopy of rim and basal foodcrusts from #3258 vessel. A-E: Images of complex rim crust with fungal micelles (My), incrusted bisaccate conifer pollen (panel B) and Streptomyces-type bacterial spores (panel E) on the surface. Damaged plant phloem and xylem microstructures, leaves surface cuticle - 50-70µk diameter broken tubes ensheathed by ca 2µk smooth wall, frazzles of thin membranes and fluffy bulk material – are designated with arrows on panels C and D, and also well recognizable on panel E. Panel F: A piece of basal crust consisting of dense homogeneous material.

## Discussion

### Emerging evidence of Endmesolithic pottery production in the region between Elbe and Oder

Coarse ceramic fragments of conical bowl #3258 and S-shaped vessel #3157 assigned to the Friesack-Boberger group is the oldest pottery found at the Friesack 4. Their morphology differs from other ceramics recovered at the site which are presented by region typical middle and late Neolithic cultures [23,24]. Similar examples of endmesolithic vessels which morphology was typologically foreign to the Mesolithic Ertebølle and the Early Neolithic Funnelbeaker pottery dominated at the area were reported earlier at the Boberg site southeast of Hamburg [54-57], and opened a question about ceramic production by endmesolithic groups in Northern Europe. Based on typology and stylistic details it was also hypothesized that these pottery might be associated with other known cultures [58,59] or considered as an evidence of adoption and flow of technologies resulted from interactions with early Neolithic settlers [40,55]. Endmesolithic ceramics of the recently described Friesack-Boberger group recovered at Friesack 4 and Rhinow 30 sites [23,24] comprise fragments of coarse pottery with certain similarity to Ertebølle and pottery from Boberg (Supplementary Fig S2). Inclusion of the well preserved shards #3258 in the group supports the notion that ceramics were manufactured by endmesolithic hunter-gatherers far south in Northern European inland, and contributes to understanding of Neolithization process in the region between Elbe and Oder. Whether the Friesack-Boberger group was developed under influence of early Neolithic cultures in Havelland reminds an open question. Although there are evidences for elder pottery in the vicinity south and north from Friesack (Linear, Stroke and Rössener ceramics [24,60,61]) no links to these groups has been found in Friesack 4 so far.

Shape, undecorated massive walls, flat base, prominent carbonized residue on the internal surface and rim crust assumed that pottery #3258 was used for cooking or stewing, set on open fire or embers. The bowl has relatively small volume and was unlikely utilized for preparation of seasonal scaled stocks or family meal but could be used for cooking of selected foods. The ware certainly belonged to a valuable household utensils: secondary puncture on the wall indicates that when the bowl got a crack it was first carefully mended and later, once probably became unrepairable, finally littered.

### Characteristics of Friesack 4 meals

*Suinae* collagen identified in foodcrust #3251.1 suggested that pork with bones, sinews or skin was cooked in this pottery. It correlates well with Neolithic archaeozoological artefacts from Friesack 4 which include boar bones [32], and corroborates the notion that this site was used as hunting station.

How carp roe was processed in the small bowl #3258? Fish roe is an ingredient of many old traditional recipes. It is considered delicacy and can be consumed grilled, fried, marinated, baked, smoked, dried, cured and also boiled in broth [62]. Proteomics did not reveal evidences of fermentation, although this food processing practice was exercised by costal settlers of the Northern Europe already in early Mesolithic [63]. No microorganisms which might have been associated with fish fermentation, as for example *Tetragenococcus halophilus*, [64-67] were among bacterial species identified in the foodcrust. Charred deposit in the interior of the bowl indicated that the food was most likely thermally processed. We hypothesized that entire carp roe sac was poached on embers in small volume of water or fumet. During cooking, the small pot was probably capped with leaves. Their charred remains encrusted in the organic deposit at the rim of the bowl are observed on electron microscopy images (Fig 5). This fish meal from Friesack is rather exquisite seasonal dish and does not belong to staple food.

It is conceivable that Friesack foods were also spiced with herbs or additionally salted. Fried-clay moulds for salt extraction (briquetage) dated on the 3^rd^ millennium BC were unearthed in Halle a.d.S. ca 140km away from Friesack 4 [68-70]. Although no proofs of saltern were found directly at Friesack, local geological prerequisites [71] and historical reports describing spontaneous crystallization of salt on soil surface in the direct proximity of the site [72] do not exclude this possibility.

### Fish meal #3258 and exploitation of aquatic resources at the Friesack 4 site

Fishing was an important part of subsistence strategy at Mesolithic Friesack. This notion is strongly supported by variety of recovered artefacts - bast knotted nets, birch-bark net-floats, fragments of dug-out canoe - and ichtioarchaeological data [27,29]. Small number of fish bones was also found in later occupational layers [32], though, no fishing or fish processing tools elucidating role of aquatic resources in transition and early Neolith were recovered at the site. Lucky finding of carp roe meal in pottery #3258 promotes the assumption that fish could be utilized as a food reserve. Fish could be caught even with simple traps during spring flooding and spawning season at Friesack site. Then, if fish was intended for processing, its entrails should be removed. Valuable roe in potter #3258, after separation from inedible offal, was cooked for immediate consume whereas the fish body might have been preserved and consumed later at other location. The assumption of delayed-return subsistence strategy might also partially explain relatively low number of fish bones in the Neolithic occupational layer representing 3% of totally 21 thousand faunal remains [32].

Proteomics applied in this work revealed details of prehistorical culinary recipes directly from charred foodcrust. This work also highlighted the impact of sample heterogeneity and environmental contaminations to the result interpretation. Being recovered from drained and cultivated peat bog, Friesack samples are heavily contaminated with remains of agricultural plants and abundant soil bacteria *Streptomyces spp.* – one of a few known bacterial genera which accumulates fats (triacylglycerols) [73,74]. Whereas proteomics distinguished between ancient and contemporary materials, common lipid- or fatty acid-based method as for example CSIA (compound specific stable isotope analysis) could be affected by presence of modern bacterial lipids in the sample.

## Acknowledgment

The authors would like to thank Drs. Sofia Traikov and Henrik Thomas from the Laboratory of Biological Mass Spectrometry and Drs. Tobias Fürstenhaupt and Michaela Wilsch-Bräuninger from the Electron Microscopy Facility at the MPI CBG in Dresden for expert technical support; Prof. Dr. Bernhard Gramsch for valuable discussion; Prof. Dr. Thomas Terberger from the Seminar for Ur- und Frühgeschichte at the Universität Göttingen for the support with ^14^C-analysis; Dr. Jonas Beran and Dr. Andrej Shevchenko for critical reading of the manuscript.

## Supporting Information

**S1 Fig.** ^14^C dating of the rim crust from pottery #3258.

**S2 Fig.** Early ceramics from Friesack 4 and Rhinow 30 tentatively attributed to Friesack-Boberger group. All ceramics except for #1 and #2 were excavated at Friesack 4; the #1 and #2 fragments – at Rhinow 30 site located 20km to the west from Friesack. Pottery fragments with inventory numbers “If” are stored in the Berlin State Museums, the remaining - in the BLDAM (Wünsdorf, Germany). Apart of ceramics shown on Fig.S2, to the Friesack-Boberger group very likely might be also attributed ceramic excavated at the Friesack site by M.Schneider [24,25]. Unfortunately these objects were lost during the Second World War. 14C-dating is shown where available. **2845/3 (Inv. No 1977:7 2845/3):** 1x body shard, coarse fired clay tempered with grit/quartz, outside light brown, inside brown-grey, slip-glaze, smooth and crackly inside, 1,0cm thick, from the lower part of a vessel with probably ca.12cm diameter. **2846.4 (Inv. No 1977:7 2846.4):** 1x rim and 3x body shards of a bowl, rounded lip, ochre-brown, outside roughened probably with stalks, inside smooth, medium coarse fired clay tempered with grit, 0.5cm thick. **2949.1 and 2 (Inv. No 1977:7 2949.1 and 1977:7 2949.2):** 1x rim and 1x body shards of jar or bowl with ca 30cm diameter, rounded lip, well smoothed surface, brown to dark-brown, medium coarse fired clay tempered with grit, 0.7cm thick. 14C dating rim shard: 4546-4366 calBC (external crust) (Poz-84049), 4857-4709 calBC (internal crust) (Poz-84050). **3038.3** (Inv. No 1977:7: 3038.3): 1x body shard, probably from a conical lower part of a vessel with ca 10cm diameter, outside grey-brown, inside and edge light brown/ochre, smooth, coarse fired clay tempered with coarse grit, 0.65-0.7cm thick, fine triangular point impressions on the outer surface. **3085.1 (Inv. No 1977:7 3085.1):** rim shard of a flat bowl with 26cm diameter, square-edged slightly rounded uneven lip, grey-brown with dark spots, medium coarse fired clay temperate with grit, 0.6-0.7cm thick. ^14^C dating: 3778-3648calBC (Poz-84051). **3127.6 (Inv. No 1977:7 3127.6):** 2x body shards of a round bowl with ca 28cm diameter, 0.6cm thick, fingertip impressions with unclear orientation, inside/outside light grey-brown. **3157.1 (Inv. No 1977:7 3157.1):** 2x body shard from shoulder of a S-shaped vessel with ca 30cm diameter, all-around impressions of vertically oriented fingertips arranged in horizontal lanes, middle coarse fired clay, tempered with grit, friable, inner black, outside brown. 14C dating: 4336-4241calBC (ARR 15048). Early Neolith? Brzesc Kujawski? **3171/1 (Inv. No 1977:7 3171/1)**: 3x body shards of a round-bodied vessel with neck/shoulder, ca 30-34cm diameter, 0.7-1.0cm thick, middle coarse fired clay tempered with grit, outer surface is structured with press- and point-impressions, porous and rough internal surface, outside grey-brown, inside black-grey. **3251.1 (Inv. No 1977:7 3251.1):** 1x body shard, shoulder of a vessel with ca 28cm diameter, 0.5-0.6cm thick, middle coarse fired clay tempered with grit, three closely placed drop-shaped impressions, smooth inner surface, grey-brown to dark-brow. 14C dating 3870+/-60 calBC. Early Neolith? Brzesc Kujawski? **3256.1 (Inv. No 1977:7 3256.1):** 1x body shard, vertically orientated fingertips/fingernails impressions arranged in horizontal lanes, coarse fired clay tempered with grit. **3258:** description in the Result section. **If 11710(4):** 1x body shard of a lower part of a vessel with ca 22-25cm diameter, regular 0.6cm drop-shaped tip-over impressions arranged in rows, 1.5cm between impressions, 1cm between rows, ochre grey-brown, smoothly roughed, middle coarse fired clay tempered with grit, 1.0cm thick. **If 11711(9):** 1xbody shard, all-around fingertips prints, temper pulled-up with nails, grey-brown, middle coarse fired clay tempered with grit, smooth inside, diameter ca 32-34cm, 0.7-0.9cm thick. **#1** (Inv.no. 1995:375/2/126): Rim, body and base shards of a vessel with S-shaped profile and conical base, decorated with 4-5mm round imprints, fired clay tempered with sand and grit, outside burnt; 24-28cm, height 29-30cm; walls 0.9cm, 1.6cm at the base. **#2** (Inv.no. 1995:375/2-169,171) 2x body shards of a S-shaped vessel, decorated with irregular imprints, fired clay tempered with sand and grit; 26cm diameter, wall 0.5-0.6cm.

**Supplementary dataset S1.** List of proteins identified in archaeological and reference samples by proteomics.

